# Organization of the Yeast Seipin Complex Reveals Differential Recruitment of Regulatory Proteins

**DOI:** 10.1101/2025.03.14.642698

**Authors:** Yoel A. Klug, Pedro Carvalho

**Author notes:** Corresponding authors: Yoel A. Klug, Pedro Carvalho.

## Abstract

Lipid droplets (LDs) are neutral lipid storage organelles that emerge from the endoplasmic reticulum (ER). Their assembly occurs in ER regions enriched with seipin which, through its homooligomeric ring-like structure, facilitates neutral lipid nucleation. In yeast, seipin (Sei1) partners with Ldb16, Ldo45 (yeast homologue of human LDAF1) and Ldo16, which regulate LD formation and consumption. How the molecular architecture of the yeast seipin complex and its interaction with regulatory proteins adapt to different metabolic conditions remains poorly understood. Here, we show that multiple Ldb16 regions contribute differently to recruiting Ldo45 and Ldo16 to the seipin complex. Using an in-vivo site-specific photo-crosslinking approach, we further show that Ldo45 resides at the center of the seipin ring both in the absence and presence of neutral lipids. Interestingly, neutral lipid synthesis leads to the recruitment of Ldo45 but not Ldo16 to the complex. Our findings suggest that the seipin complex serves as a pre-assembled scaffold for lipid storage that can be remodeled in response to increased neutral lipid availability.

## Introduction

Lipid droplets (LDs) are ubiquitous storage organelles for neutral lipids, such as diacylglycerol (DAG), triacylglycerol (TAG) and steryl esters (SE), with a central role in lipid and energy metabolism (Olzmann and Carvalho, 2019; Walther and Farese, 2012). LD formation depends on neutral lipid synthesis (Sandager et al., 2002) and occurs at the endoplasmic reticulum (ER) (Jacquier et al., 2011; Kassan et al., 2013). Neutral lipids are synthesized within the ER and at low concentrations disperse within the bilayer. At a critical concentration (5-10%), neutral lipids phase-separate and coalesce, forming a lens-like structure within the ER leaflets. As the concentration of neutral lipids rises, the lens expands and gives rise to a nascent LD, which buds out of the ER towards the cytosol (Chorlay et al., 2019). This process results in the LD’s unique architecture, comprised of a neutral lipid core encapsulated by an ER derived phospholipid monolayer (Fujimoto and Parton, 2011; Tauchi-Sato et al., 2002) and a set of LD specific proteins (Bersuker et al., 2018; Currie et al., 2014).

Seipin, an evolutionary conserved ER membrane protein localizes to sites of LD biogenesis and has emerged as the main LD assembly factor(Fei et al., 2008; Grippa et al., 2015; Salo et al., 2016; Wang et al., 2014; Wang et al., 2016). In humans, seipin is encoded by the Bernadelli-Seip congenital lipodystrophy type 2 (BSCL2) gene, often mutated in familial forms of lipodystrophies(Magré et al., 2001). At the cellular level, loss of seipin results in aberrant LD morphology and numbers, with many small and clustered LDs and a few super-sized ones(Boutet et al., 2009; Fei et al., 2008; Szymanski et al., 2007). Aside from defects in LD biogenesis, seipin mutations also prevent correct ER-LD contacts(Grippa et al., 2015; Salo et al., 2016), LD maturation(Wang et al., 2016) and maintenance (Salo et al., 2019).

In *S. cerevisiae* (hereon termed yeast), seipin (Sei1) function requires its binding to Ldb16(Grippa et al., 2015; Wang et al., 2014), an ER membrane protein specific to certain fungi. LD defects observed in *sei1Δ*, *ldb16Δ* and *sei1Δldb16Δ* cells are indistinguishable (Han et al., 2015; Wang et al., 2014) and can be rescued by human seipin (Wang et al., 2014), suggesting that in yeast, seipin function is distributed between Sei1 and Ldb16.

Seipin has two transmembrane segments at the N and C termini with an ER luminal loop between them (Lundin et al., 2006) and forms a homooligomer (Binns et al., 2010). Cryogenic electron microscopy (Cryo-EM) studies of seipin from human (Li et al., 2024; Yan et al., 2018), fly (Sui et al., 2018) and yeast (Arlt et al., 2022; Klug et al., 2021) show that seipins form homooligomeric rings of 11 (Yan et al., 2018), 12 (Sui et al., 2018) and 10 (Arlt et al., 2022; Klug et al., 2021) subunits, respectively. A recent study showed that human seipin, in complex with the adipose specific protein adipogenin, can adopt a 12 subunit conformation (Li et al., 2024), though the physiological relevance of having two distinct seipin conformations is still unclear. Regardless of the number of subunits, oligomerization into a ring-like structure is essential for seipin function (Sui et al., 2018; Yan et al., 2018). At the single protomer level, all seipins adopt a similar structure: an 8-strand β-sandwich in their luminal domain (Arlt et al., 2022; Klug et al., 2021; Li et al., 2024; Sui et al., 2018; Yan et al., 2018). In human and fly seipins the luminal domain is capped with a hydrophobic helix, which in the oligomer resides at the center of the seipin ring and protrudes into the ER bilayer (Sui et al., 2018; Yan et al., 2018). Molecular dynamic (MD) simulations backed by mutagenesis suggest that two serine residues in the hydrophobic helix bind neutral lipids via their hydroxyl moiety, thus aiding in TAG nucleation and subsequent LD formation (Prasanna et al., 2021; Renne et al., 2022; Zoni et al., 2021). However, in yeast, instead of a hydrophobic helix, seipin has two short hydrophilic helices that cannot enter the ER bilayer (Arlt et al., 2022; Klug et al., 2021). In support of this, we have previously shown via photo-crosslinking that Ldb16 resides at the center of the Sei1 ring (Klug et al., 2021) and likely serves as the neutral lipid binding entity of the yeast seipin complex using similar chemistry as human and fly seipin, to bind neutral lipids such as TAG and SE (Klug et al., 2021; Klug et al., 2024; Renne et al., 2022). Overall, these findings suggest a unifying mechanism for LD formation by seipins (Klug et al., 2024).

Apart from binding neutral lipids, the hydrophobic helix in human seipin was shown to bind to Lipid droplet assembly factor 1 (LDAF1)/promethin (Castro et al., 2019; Chung et al., 2019; Prasanna et al., 2021), a conserved ER membrane protein homologous to yeast Ldo45 (Castro et al., 2019; Eisenberg-Bord et al., 2018; Teixeira et al., 2018) and Fly CG32803 (Chartschenko et al., 2021). In yeast, Ldo45 is encoded together with Ldo16 by a consecutive, partly overlapping, open reading frame and is generated by alternative splicing with Ldo16 included in the Ldo45 sequence (Eisenberg-Bord et al., 2018; Teixeira et al., 2018). Like Ldo45, Ldo16 binds the yeast seipin complex but has a distinct role. While Ldo45 regulates LD morphology and TAG accumulation (Eisenberg-Bord et al., 2018; Teixeira et al., 2018), similarly to LDAF1 (Chung et al., 2019), Ldo16 plays a role in lipophagy (Álvarez-Guerra et al., 2024; Diep et al., 2024; Teixeira et al., 2018).

While the precise function of LDAF1 and Ldo45 remains poorly defined, these proteins likely play a regulatory role. Consistent with this, deletion of LDAF1 or Ldo45 results in a milder LD phenotype than seipin mutations (Chung et al., 2019; Eisenberg-Bord et al., 2018; Teixeira et al., 2018).

Here we used a previously established photo-crosslinking approach (Klug et al., 2021) to investigate the organization of seipin complex, focusing on the Sei1 interactors Ldb16, Ldo45 and Ldo16. Combining this methodology with an LD induction assay revealed that Ldo45 occupies the central position within the Sei1-Ldb16 complex, nestled close to Ldb16, regardless of the presence of neutral lipids. Moreover, we observed that Ldo45 and Ldo16 interact with distinct regions of Ldb16 and that their expression is modulated by neutral lipid levels. Thus, in yeast the seipin complex assembles prior to LD formation and retains this organization throughout the process. Moreover, Ldb16 emerges as a key player in the yeast seipin complex, not only binding neutral lipids but also directly anchoring regulatory proteins like Ldo45 at its core, thus orchestrating lipid droplet formation.

## Results

### Ldo45 is positioned in the center of the Sei1 ring adjacent to Ldb16

We previously used a site-specific photo-crosslinking strategy to map the interactions among the protomers of the Sei1 ring (Klug et al., 2021). This approach also allowed us to map Sei1 interaction with its partner Ldb16. This photo-crosslinking system employs an amber codon suppressor tRNA and a modified tRNA synthetase to incorporate a photoreactive amino acid derivative, benzoyl-phenylalanine (Bpa), at sites designated by an amber stop codon. In cells expressing this system and grown in the presence of Bpa, UV irradiation induces crosslinking between Bpa-labeled probes and nearby proteins (Chin et al., 2003) (Figure 1A).

**Figure 1.**
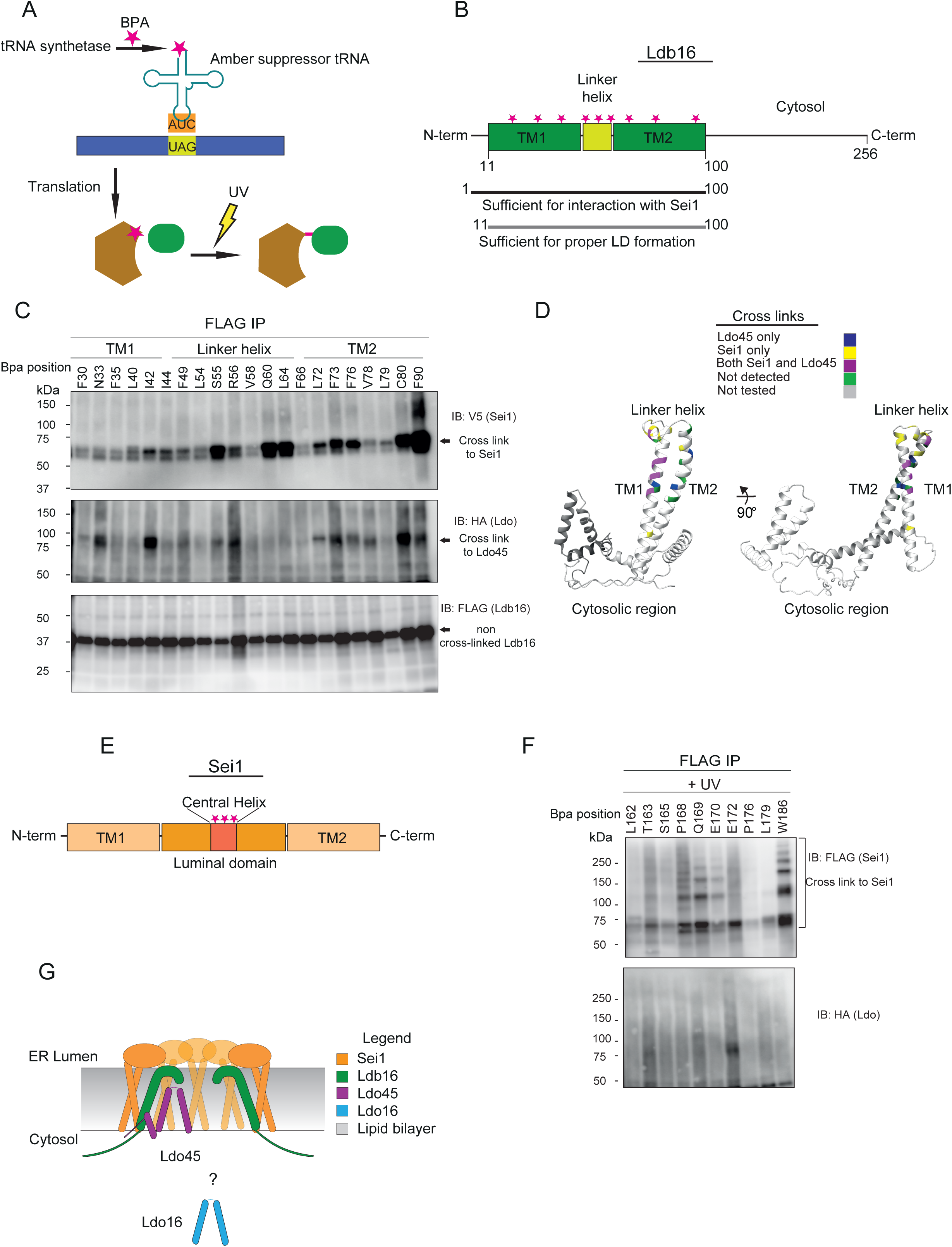
A) Site specific photo-crosslinking method used in this study. B) Schematic of Ldb16 domains, functional regions and Bpa positioning. C) Ldb16Δ cells expressing endogenously HA-tagged Ldo, V5-tagged Sei1 and plasmid-borne Ldb16-FLAG with a photoreactive Bpa at the indicated positions were subjected to UV irradiation. Solubilized membranes were subjected to immunoprecipitation with anti-FLAG antibodies, and bound proteins were analyzed by immunoblotting with FLAG, V5 and HA antibodies. D) Schematic representation of the Ldb16-Ldb16, Ldb16-Sei1 and Ldb16-Ldo site-specific photo-crosslinks obtained Figure 1C and S1B. E) Schematic of Sei1 domains and Bpa positioning. F) Sei1Δ cells expressing endogenously HA-tagged Ldo, and plasmid-borne Sei1-FLAG with a photoreactive Bpa at the indicated positions were subjected to UV irradiation. Solubilized membranes were subjected to immunoprecipitation with anti-FLAG antibodies, and bound proteins were analyzed by immunoblotting with FLAG and HA antibodies. G) Sei1, Ldb16 and Ldo organization in the ER bilayer.

We employed the same approach to investigate the interactions between Ldb16 and the additional components of the seipin complex, namely Sei1, Ldo45 and Ldo16. Ldb16 has an N-terminal membrane domain, consisting of two TMs with a short linker helix between them shown to be important for proper LD formation (Klug et al., 2021), followed by a large C-terminal region in the cytosol (Figure 1B). Ldb16 truncations containing only the N-terminal membrane domain (residues 1-100) support both Sei1 binding and proper LD formation (Figure 1B) (Wang et al., 2014). We generated Ldb16 variants with individual Bpa probes along the N-terminal membrane domain, including transmembrane domain 1, linker helix and transmembrane domain 2 (Figure 1B, 1C and S1B). These variants we expressed in cells where the other subunits of the Sei1 complex were epitope tagged to facilitate detection and expressed their endogenous loci. Sei1 was expressed as a V5 fusion while Ldo16 and Ldo45, partly encoded by the same gene (Eisenberg-Bord et al., 2018)(Teixeira et al., 2018), were expressed as HA fusion proteins (Figure S1A). Several Bpa probes in Ldb16 TM1, linker helix and TM2 crosslinked to Sei1-V5 (Figure 1C and D and S1B). These crosslinks, detected after immunoprecipitation of Ldb16, were highly specific and detected only in UV-irradiated cells at defined Bpa positions (Figure 1C and D and S1B). Importantly, they are consistent with our structural model showing that Ldb16 is positioned within the homodecameric Sei1 ring (Klug et al., 2021).

We also observed crosslinks between Ldb16 and Ldo45. When probes were introduced within TM1, linker helix and TM2 of Ldb16, crosslinks to Ldo45 were detected (Figure 1C and D and S1B). Ldo45 crosslinks were particularly strong for Bpa probes inserted at the Ldb16 linker helix (S55 and R56), TM1 (N33 and I42) and TM2 (F72, F73 and C80). While Ldo16 is the most abundant between the two Ldo proteins and co-precipitates with other components of the Sei1 complex (Teixeira et al., 2018), no crosslinks were detected between Ldb16 and Ldo16.

Human seipin has been shown to bind LDAF1, the human homolog of Ldo45, via its hydrophobic central helix. To further examine Ldo45 and Ldo16 positioning in the complex, we repeated the photo-crosslinking experiment but this time with the Bpa probe situated in the Sei1 central helix (Figure 1E), and Ldo45 and Ldo16 expressed as HA fusion proteins as before. Following UV irradiation and Sei1 immunoprecipitation we detected Sei1-Sei1 crosslinks (Figure 1F and Figure S1C), consistent with what we have previously reported (Klug et al., 2021). However, no crosslinks between Sei1 and Ldo45 or Ldo16 were observed (Figure 1F and Figure S1C). To exclude that only a minor fraction of Ldo45 was crosslinking, we enriched Ldo proteins by immunoprecipitation, but no crosslinks were detected even under these conditions (Figure S1D). Thus, Ldo proteins do not appear to bind directly to Sei1 and are recruited to the seipin complex via interactions with Ldb16. Moreover, these data suggest that Ldb16 adopts a pivotal position within the seipin complex with its membrane domain interacting both with Sei1 and Ldo45 but not Ldo16 (Figure 1G).

### Ldb16 interacts with Ldo45 and Ldo16 through distinct regions

Consistent with our crosslinking data, previous immunoprecipitation experiments showed that Ldb16 is required for the interaction between Sei1 and Ldo proteins (Teixeira et al., 2018). Our crosslinking experiments showed direct interactions between the Ldb16 linker helix and Ldo45 (Figure 1C). This is consistent with co-immunoprecipitation experiments in mammalian cells, which showed that the intramembrane hydrophobic helix of seipin, equivalent to the Ldb16 linker helix, is required for interaction with the Ldo45 homologue LDAF1 (Chung et al., 2019). We used co-immunoprecipitation to test if other Ldb16 regions contribute to the interaction with Ldo45 and Ldo16.

We observed that mutation (Ldb16 6A) or deletion (Ldb16 ΔH) (Figure 2A and B) of the Ldb16 linker helix showed reduced binding to both Ldo45 and Ldo16 (Figure 2C and D). Next, we analyzed the contribution of Ldb16 cytosolic domain by generating (1-133), a truncation consisting mostly of the membrane domain (lacking residues 134-256) (Figure 2A). This mutant allows normal LD formation (Grippa, 2016) and binds efficiently to Sei1 but shows reduced interaction with both Ldo45 and Ldo16 (Figure 2C). Interestingly, when the truncation of the cytosolic domain was combined with mutations in linker helix in mutants (1-133 6A) and (1-133 ΔH) (Figure 2A and B) we observed a further reduction of the interaction with both Ldo proteins (Figure 2C and D). Ldo16 exhibited the strongest reduction under these conditions (Figure 2C and D). Noteworthy, the linker helix mutants 1-133 6A and 1-133 ΔH showed close to 50% lower levels of Ldo co-immunoprecipitation when compared to 1-133 (Figure 2E). These results suggest that both the linker helix and the cytosolic region of Ldb16 are involved in Ldo protein recruitment.

**Figure 2.**
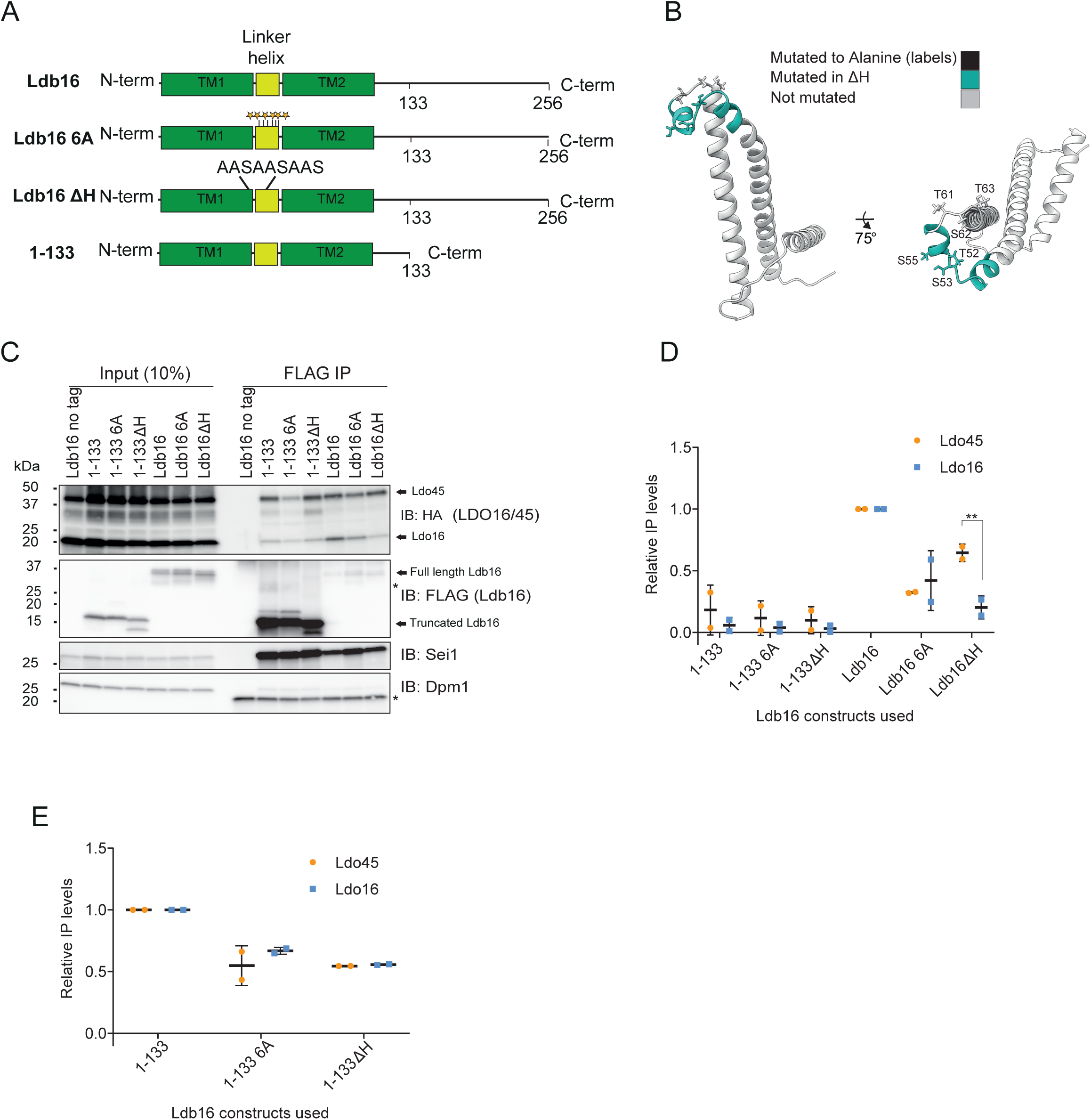
A) Schematic representation of the Ldb16 mutants used. B) *In-silico* structural prediction by trRosetta of residues 1-133 of Ldb16. Mutations made are highlighted and labelled. C) Genomic expressed Ldb16-FLAG constructs were immunoprecipitated from cells expressing endogenously HA-tagged Ldo using FLAG beads from membranes solubilized with 1% GDN. Eluted proteins were analyzed by SDS–PAGE and immunoblotting. Dpm1 was used as a loading control. *=Non specific bands. D) Quantification of Ldo IP levels relative to cells expressing WT Ldb16. Ldo levels were normalized to input and Ldb16 expression levels. n=2. Difference was tested using a two-sided Fischer’s T-test. Error bars are S.D. (**p < 0.01). E) Quantification of Ldo IP levels relative to cells expressing Ldb16 1-133. Ldo levels were normalized to input and Ldb16 expression levels. n=2. Error bars are S.D.

### Pre-assembled seipin complex is stable during LD biogenesis

LDs are highly dynamic organelles, continuously being assembled and consumed according to cellular metabolism. It remains unknown if there are changes in the composition and/or organization of the seipin complex during the LD life cycle. To address this question, we used immunoprecipitation to compare the composition of the seipin complex in wild-type (WT) and Δ4 mutant cells, which are devoid of LDs due to mutations in the four neutral lipid synthesizing enzymes: the TAG acyltransferases Dga1 and Lro1, and the SE acyltransferases Are1 and Are2. Although seipin complex components appeared slightly reduced in Δ4 cells, Sei1 and Ldo45 co-precipitated with Ldb16 irrespective of the presence of LDs (Figure 3A), as expected (Wang et al., 2024). Interestingly, a much lower amount of Ldo16-HA co-precipitated with Ldb16 in Δ4 cells (Figure 3A).

**Figure 3.**
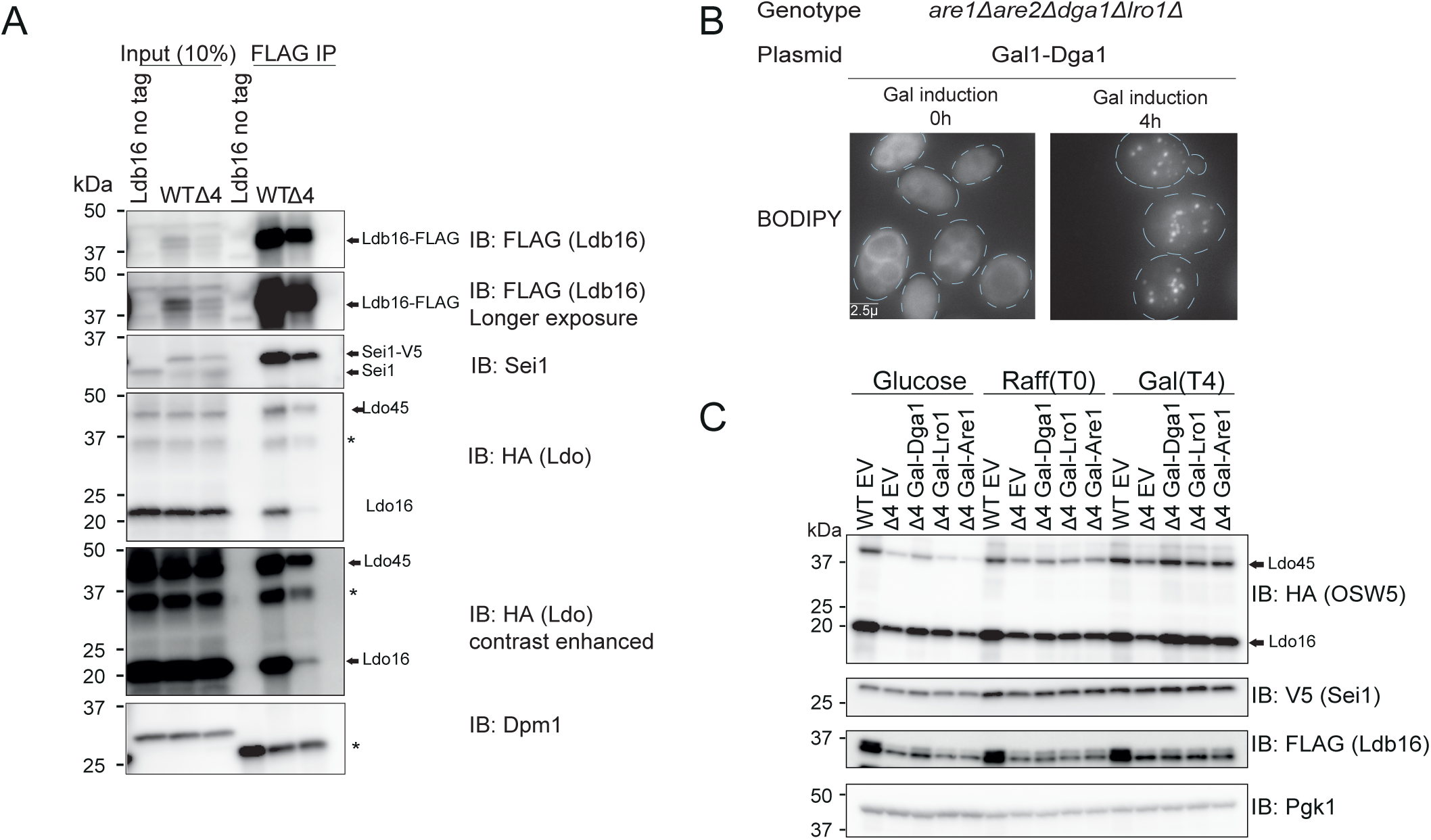
A) Plasmid-borne WT Ldb16-FLAG was immunoprecipitated from WT cells or Δ4 cells expressing endogenously HA-tagged Ldo and V5-tagged Sei1 using FLAG beads from membranes solubilized with 1% GDN. Eluted proteins were analyzed by SDS–PAGE and immunoblotting. Dpm1 was used as a loading control. * denotes non-specific bands. B) Analysis of LDs in cells with the indicated genotype after staining with the neutral lipid dye BODIPY 493/503. Lipid droplet biogenesis was induced by the addition of 2% galactose (Gal) to cells with indicted genotype expressing DGA1 from the GAL1 promoter. Time is in hours. Scale bar corresponds to 2.5μm. C) Δ4 Ldb16Δ cells expressing endogenously HA-tagged Ldo, V5-tagged Sei1 and plasmid-borne Ldb16-FLAG and denoted acyl-transferase under a galactose promoter were grown in 2% glucose, then 2% raffinose (Raff) and lastly in 2% galactose (Gal) for four hours. Cell lysates were analyzed by immunoblotting with FLAG, V5 and HA antibodies. Pgk1 was used as a loading control.

To further analyze the seipin complex during LD formation, we tested if the levels of its subunits were affected by the presence of neutral lipids. To this end, plasmids encoding individual neutral lipid synthesizing enzymes expressed from the conditional galactose-inducible promoter were introduced into Δ4 cells. When grown in the absence of galactose, these cells were devoid of LDs, with galactose addition resulting in synchronous LD formation as shown in Figure 3B for Dga1 expression. Consistent with our earlier observation, steady state levels of Ldb16 and Ldo proteins were reduced in the Δ4 mutant in comparison to WT cells while Sei1 was present at comparable levels in both cells (Figure 3C). Induction of neutral lipid synthesis for 4 hours increased the steady state levels of Ldb16, Ldo45 and Ldo16. Interestingly, this increase was observed upon induction of either TAG or SE (Figure 3C), suggesting that it results from the formation of LDs rather than a specific type of neutral lipid.

Next, we combined this LD biogenesis assay with our photo-crosslinking approach (Figure 4A), which is highly sensitive in capturing position-specific interactions, to search for potential changes in conformation and/or organization within the seipin complex. Upon introducing the photo-crosslinking system and the Ldb16 plasmids with individual Bpa probes at chosen positions, we analyzed interactions within the seipin complex before (T0) and four hours after (T4) the induction of LD biogenesis (Figure 4A and B). Irrespective of the absence or presence of LDs, Ldb16 showed robust crosslinks to both Sei1 and Ldo45 (Figure 4B and C). Importantly, the stronger Ldb16 crosslinks were maintained between the two conditions suggesting that the relative position of Ldb16, Sei1 and Ldo45 membrane domains was similar in the absence and presence of LDs (Figure 4B and C). Interestingly, the strength of most crosslinks was increased upon LD induction (Figure 4D), likely due to the higher steady state levels of Ldb16 and Ldo45 under these conditions.

**Figure 4.**
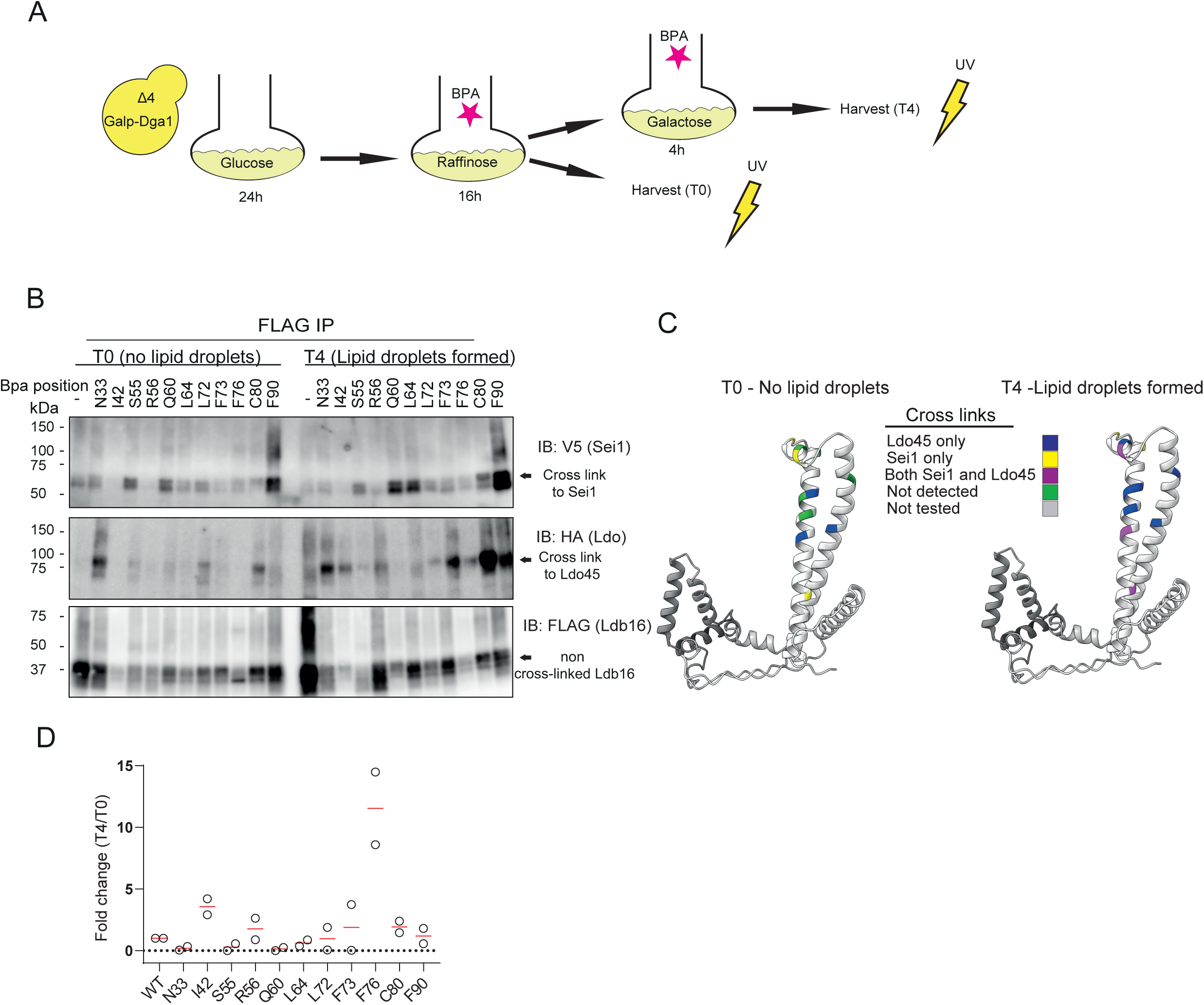
A) Lipid droplet induction method coupled with site specific photo-crosslinking. B) Δ4 Ldb16Δ cells expressing endogenously HA-tagged Ldo, V5-tagged Sei1 and plasmid-borne Ldb16-FLAG with a photoreactive Bpa at the indicated positions, were subjected to UV irradiation before (T0) and four hours after (T4) the addition of 2% galactose. Solubilized membranes were subjected to immunoprecipitation with anti-FLAG antibodies, and bound proteins were analyzed by immunoblotting with FLAG, V5 and HA antibodies. C) Schematic representation of the Ldb16-Ldb16, Ldb16-Sei1 and Ldb16-Ldo site-specific photo-crosslinks obtained in (B). D) Quantification of photo-crosslink enrichment at T4 compared to T0 shown in fold change. Crosslink intensities from each time point were normalized to the corresponding WT band.

**Figure 5.**
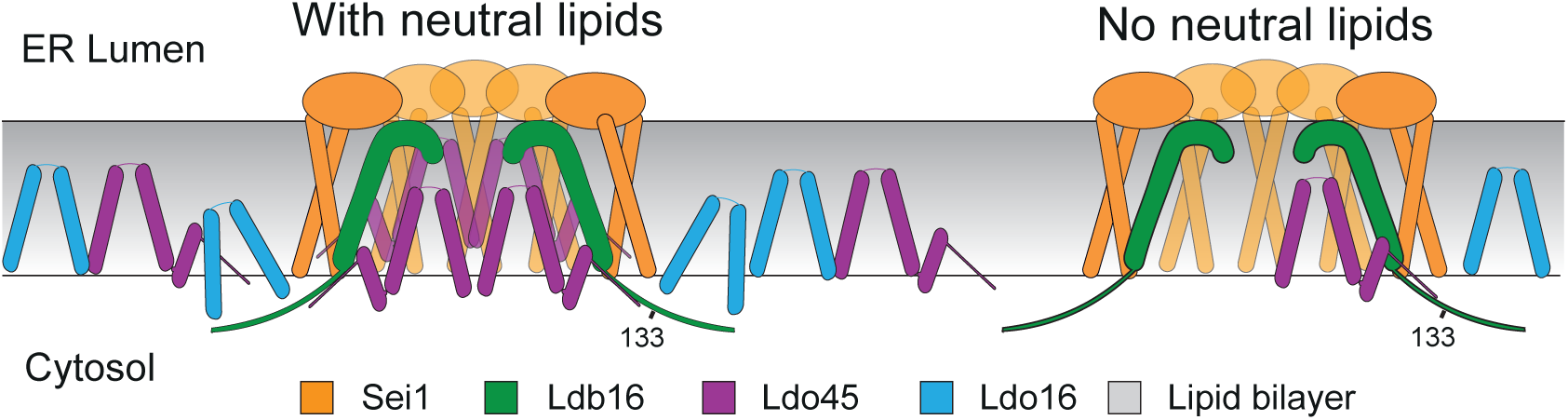
Model of Seipin complex assembly. Regardless of neutral lipid abundance, the seipin complex organizes with Ldb16 (green) and Ldo45 (purple) at the centre of the seipin ring (orange) with Ldo16 (blue) sitting outside the ring. Ldo proteins are recruited by Ldb16, with Ldo45 adjacent to the Ldb16 linker helix and both Ldo45 and Ldo16 binding the cytosolic region of Ldb16. When neutral lipids accumulate within the seipin ring, Ldo protein levels rise and they are retained at the complex.

## Discussion

In this study, we mapped the molecular organization of the yeast seipin complex and characterized its changes during lipid droplet (LD) formation.

We demonstrated that pre-assembled seipin complexes are present in cells even in the absence of LDs. Using site-specific photo-crosslinking, we observed that Ldb16 interacts directly with Ldo45, positioning it at the center of the seipin ring, while Ldo16 appears to distribute more peripherally. Interestingly, the levels of both Ldo45 and Ldo16 increase with neutral lipid synthesis, consistent with their regulatory function on the seipin complex.

Our *in-vivo* site-specific photo-crosslinking analysis of the seipin complex revealed that Ldb16 functions as a linchpin, bringing together the various subunits of the complex. The Ldb16 membrane domain plays a key role in these interactions consistent with it being the minimal region required for normal LD formation (Grippa, 2016; Wang et al., 2014). The interactions between Ldb16 and Sei1 agree with our previous photo-crosslinking analysis, where the photoreactive Bpa probes were introduced in Sei1 (Klug et al., 2021) and further support the idea that the Sei1 ring scaffolds and positions Ldb16 for neutral lipid binding via its linker helix.

Among the various crosslinks to Ldo45, there are two residues within the Ldb16 linker helix that were shown to be important to concentrate neutral lipids for LD formation (Klug et al., 2021; Renne et al., 2022). Previous studies showed that the equivalent helix in human seipin interacted with LDAF1 (Chung et al., 2019; Prasanna et al., 2021) further suggesting that Ldo45 and LDAF1 may use similar mechanisms to regulate the function of seipin complexes in yeast and human cells, respectively. Our systematic crosslinking analysis did not detect crosslinks between either Sei1 or Ldb16 with Ldo16, despite this being the most abundant of the Ldo proteins. On the other hand, robust and reproducible interactions between Ldo16 and the core subunits of the seipin complex, Sei1 and Ldb16, are detected by immunoprecipitation. In cells expressing the Ldb16 truncation mutant (1-133) these interactions are strongly diminished suggesting Ldb16 cytosolic domain is important to stabilize the interactions with the Ldo proteins.

We detected this general organization of the yeast seipin complex both in cells with and without LDs consistent with recent immunoprecipitation experiments (Wang et al., 2024). Interestingly, the levels of both Ldo proteins increase with neutral lipid synthesis while the levels of both Sei1 and Ldb16 remain largely unchanged. Concomitant with the higher Ldo45 levels upon neutral lipid synthesis we detected stronger crosslinks with Ldb16. This suggests that that this condition increases Ldo45 stoichiometry and/or affinity to the seipin complex. In the future it will be important to dissect the functional consequences of these changes. However, considering that Ldo45 interacts with the seipin complex directly at the region required for neutral lipid concentration, it is appealing to speculate that it may modulate, and perhaps enhance, that activity. Irrespective of the mechanism, our findings support the notion that Ldo45 is important for early steps of LD formation as previously proposed (Eisenberg-Bord et al., 2018; Teixeira et al., 2018) We previously showed that the expression of Ldo16 and Ldo45 was independently regulated and differentially affected by cellular metabolism (Teixeira et al., 2018). How these proteins are regulated upon activation of neutral lipid synthesis should also be investigated in the future.

Overall, our data identify Ldb16 as the major hub in the yeast seipin complex for protein recruitment and neutral lipid binding and collectively suggest a framework for how the yeast seipin complex assembles throughout LD formation.

## Materials and Methods

### Reagents

BODIPY493/503 was purchased from Invitrogen. Antibodies used in this study were anti-FLAG M2-Peroxidase (HRP), Clone M2-A8592 (Sigma Aldrich) product number A8592, anti-HA High affinity (clone 3F10) product number 11867431001 (Roche), PGK1 Monoclonal (22C5D8) product number 459250 (Invitrogen), DPM1 monoclonal (5C5A7) product number A6429 (Thermo Fisher Scientific) and V5 monoclonal (D3H8Q) product number 13202 (Cell Signalling). Polyclonal anti-Sei1 (rabbit), anti-Ldb16 (rabbit) antibodies were previously described (Teixeira et al., 2018).

### Yeast strains and Plasmids

The strains used are isogenic either to BY4741 (*MATa ura3Δ0 his3Δ1 leu2Δ0 met15Δ0*) or FY251 (*MATa ura3-52 his3Δ200 leu2Δ1 trp1Δ63*) and are listed in the Table S1. Tagging of proteins and individual gene deletions were performed by standard PCR-based homologous recombination (Longtine et al., 1998) and standard yeast molecular genetics protocols(Guthrie and Fink, 1991). Multiple gene deletion strains were performed by CRISPR-based gene editing (adapted from (Laughery et al., 2015)). Briefly, a single guide RNA sequence targeting the desired region of the gene of interest was designed using the online software Benchling (Biology Software, 2021) and http://wyrickbioinfo2.smb.wsu.edu. The single guide RNA was cloned into the pML107 vector (Laughery et al., 2015), containing a Cas9 endonuclease from *Streptococcus pyogenes*. This plasmid, along with a PCR-amplified template containing the desired modification, was transformed using standard yeast transformation protocol.

Plasmids used are based on pRS316, pRS415, pRS416 and pRS425 (Christianson et al., 1992; Sikorski and Hieter, 1989) and listed in Table S2.

Galp-Dga1, Galp-Lro1and Galp-Are1 were expressed using a galactose promoter followed by a cyc1 terminator. 3xFLAG-tagged Sei1 was expressed from the native Sei1 promotor (494 bp upstream of the *SEI1* ORF) or an *ADH1* promotor, and followed by the *ADH1* terminator. 3xFLAG-tagged Ldb16 was expressed from the *ADH1* promoter followed by the *ADH1*-terminator. The oligonucleotides used for generation of strains and plasmids can be found in Table S3.

### Culture conditions

Cells were cultured in synthetic defined glucose media (SD), unless otherwise indicated. SD contained per liter: 6.7 g yeast nitrogen base with ammonium sulfate (YNB; MP biomedicals), 0.6 g complet esupplement mixture without histidine, leucine, tryptophan and uracil (CSM-HIS-LEU-TRP-URA; MP biomedicals), and 20 g glucose (Sigma-Aldrich). Media was supplemented with histidine (60 µM), leucine (1.68 mM), uracil (0.2 mM) and tryptophan (0.4 mM) as required.

All cultures were incubated at 30°C with shaking at 200 rpm. Culture density was determined by measuring turbidity at 600 nm (OD600) using a GENESYS 10S UV-VIS spectrophotometer (Thermo Scientific).

### *In vivo* site-specific crosslinking

Site specific crosslinking was conducted as previously described(Carvalho et al., 2010). Briefly, *Ldb16Δ* or *Ldb16Δare1Δare2Δdga1Δlro1Δ* with *sei1-V5 and OSW5-HA* cells were transformed with two plasmids, one encoding both for a modified tRNA synthetase capable of charging the unnatural amino acid benzoyl phenylalanine (BPA) on a tRNA as well as amber stop codon suppressor tRNA, and a second plasmid encoding ADH1-promotor expressed Ldb16-3xFLAG with individual amber codons. For *Ldb16Δ* based strains, cells carrying both plasmids were pre-cultured in SD for 8 hours, transferred to 100 mL of the same media supplemented with BPA to a final concentration of 0.3 mM (from a 0.3M in 1M NaOH freshly prepared stock) and cultured to mid-exponential phase (OD ∼ 1).

For *Ldb16Δare1Δare2Δdga1Δlro1Δ* based strains, cells carrying both plasmids were pre-cultured in SD for 8 hours then diluted and grown for 24h in SD till they reached OD1. Next, cells were diluted to OD 0.25 into 100ml synthetic defined raffinose media (as SD but with 2% raffinose as carbon source) supplemented with BPA to a final concentration of 0.3 mM (from a 0.3M in 1M NaOH freshly prepared stock). Cells were left for 16 hours at 30C° reaching to mid-exponential phase (OD ∼ 1).

Cells were spun down and washed with water and 100ml synthetic defined galactose media was added (as SD but with 2% galactose as carbon source) supplemented with BPA to a final concentration of 0.3 mM to induce *DGA1* expression and left to grow for 4 hours.

Cells were harvested by a centrifugation for 2 min at 3000 g and resuspended in 2 mL of cold water. Half of the cells were transferred to a 12 well plate and subjected to UV irradiation for 1 hour at 4°C using a B-100AP lamp (UVP, CA). The other half of the cells was incubated on ice and served as non-irradiated control. After UV irradiation, cells were harvested by centrifugation for 2 min at 3000 g. Both irradiated and control cells were lysed in lysis buffer (50mM Tris.HCl [pH7.4], 200 mM NaCl, 1mM EDTA, 1 mM PMSF (Roche) and 1x cOmplete protease inhibitor cocktail (Roche)by 5-6 × 1 min cycles of bead beating. Lysates were cleared by a 10 min centrifugation at 600g. Cleared lysates were centrifuged at 100,000 g (25 min at 4°C) in an Optima Max Tabletop Ultracentrifuge in a TLA 45 rotor (Beckman Coulter) to obtain crude membrane fractions. The membrane pellet was resuspended in denaturing buffer (50mM Tris.HCl [pH7.4], 1mM EDTA, 1% SDS, 2M urea) and solubilized at 65°C for 30-40 min with vigorous shaking. Insolubilized material was pelleted by centrifugation (15 min, 13000 g). The solubilized material was diluted with lysis buffer supplemented with 1% Nonidet P-40 and incubated overnight with anti-FLAG M2 magnetic beads – m8823 (Sigma-Aldrich). Beads were washed 3 times with lysis buffer/1% Nonidet P-40 and bound proteins eluted with SDS buffer and analyzed by immunoblotting.

### Native Immunoprecipitation

Cells were in SD until mid-exponential phase (OD ∼1). Cells corresponding to 50 OD were then harvested by centrifugation at 3000 g and washed with lysis buffer (50 mM Tris.HCl [pH7.4], 200 mM NaCl, 1 mM EDTA, 2 mM phenylmethylsulfonyl fluoride and 1x cOmplete protease inhibitor cocktail). Lysates and crude membrane fractions were prepared as described above. Detergent extracts were prepared by solubilizing crude membrane fractions in lysis buffer/1% glyco-diosgenin (GDN) (GDN101 anatrace). Insolubilized material was cleared by centrifugation (20,000 g, 15 min). The cleared detergent extracts were incubated overnight at 4°C with FLAG M2 magnetic beads – m8823 (Sigma-Aldrich). Beads were washed 3 times with lysis buffer/0.015% GDN, eluted with SDS-PAGE sample buffer and analyzed by immunoblotting. The input corresponds to 10% of the total extract used for IP.

### Gel electrophoresis and immunoblotting

For protein quantification of whole cell lysates, cells were lysed using NaOH as described previously (Kushnirov, 2000). Briefly, cells pellets corresponding to 1 OD were suspended in 0.15 M NaOH and incubated on ice for 10 minutes. Cells were pelleted, and resuspended in Laemmli sample buffer (Laemmli, 1970) and incubated 65°C for 10 minutes with vigorous shaking. Debris was pelleted by a short spin, and samples were loaded on a 4-20% gradient SDS-polyacrylamide gel, separated by electrophoresis and blotted to a PVDF membrane.

### Fluorescence microscopy analysis of LDs

For microscopy analysis of LDs, *Ldb16Δare1Δare2Δdga1Δlro1Δsei1-VS,OSW5-HA* cells were grown as described above for the site specific photo crosslinking under galactose induction. Cells were collected before and 4 hours after galactose induction Lipid droplets were visualised with BODIPY 493/503 (1 µg/mL).

Fluorescence microscopy was performed using a Zeiss Axio Observer.Z1 equiped with a Hamamatsu Orca Flash 4.0 digital CMOS camera. Images were acquired using an A Plan-APOCHROMAT 100x objective (*N.A.* 1.4). BODIPY fluorescence was analysed using a GFP-fluorescence set up consisting of a 485/20 bandpass excitation-filter (Zeiss), a 410/504/582/669-Di01 quad dichroic mirror, and a 525/30 bandpass emission filter. Microscope was controlled using Slidebook 6.0 software (3i).

### Statistics and reproducibility

Statistical significance and p-values were calculated in GraphPad Prism 7 using paired t-testing (2-tailed). Graphs were plotted in Prism. The following figure panels show representative data from at least three independent biological replicates that showed similar results: Figs 1C and 3C and extended data Figs S1B. The following figure panels show representative data from at least two independent biological replicates that showed similar results: Figs 2C, 2D, 2E, 3A, 4B and 4D. The following figure panels show a single experiment: Figs 1F and extended data Figs S1A, S1C and S1D.

## Acknowledgments

We thank M. V. D. Weijer, N. Sergejevs and T. Williams for critical reading of this manuscript. Y. A. Klug and P. Carvalho were supported by a BBSRC grant (BB/W015722/1).

## Author contributions

Y. A. Klug performed the experiments and data analysis. Y. A. Klug and P. Carvalho conceived the study and wrote the manuscript.

