## Supplementary data for "Organization of the Yeast Seipin Complex Reveals Differential Recruitment of Regulatory Proteins"

### Figure S1

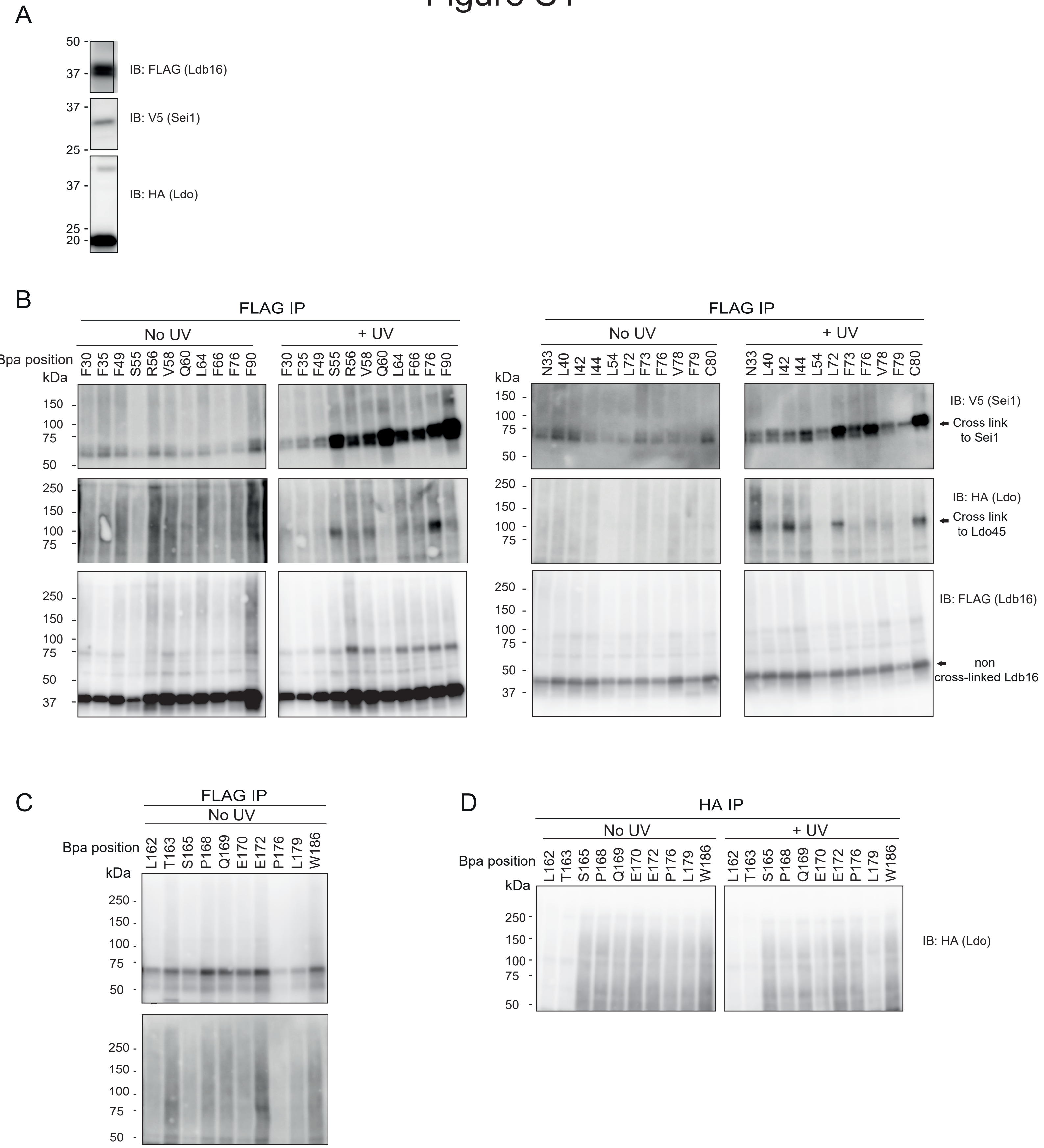
A
IB: FLAG (Ldb16)
IB: V5 (Sei1)
IB: HA (Ldo)
B
FLAG IP
No UV
+ UV
Bpa position
kDa
F30 F35 F49 S55 R56 V58 Q60 L64 F66 F76 F90
N33 L40 I42 I44 L54 L72 F73 F76 V78 F79 C80
IB: V5 (Sei1)
Cross link to Sei1
IB: HA (Ldo)
Cross link to Ldo45
IB: FLAG (Ldb16)
non cross-linked Ldb16
C
FLAG IP
No UV
Bpa position
kDa
L162 T163 S165 P168 Q169 E170 E172 P176 L179 W186
D
HA IP
No UV
+ UV
Bpa position
kDa
L162 T163 S165 P168 Q169 E170 E172 P176 L179 W186
IB: HA (Ldo)

Figure S1

A) Western blot of tagged proteins at steady state that were used in the photo-crosslinking experiments.
B) Full western blots of Figure 1C shown with and without UV. Ldb16Δ cells expressing endogenously HA-tagged Ldo, V5-tagged Sei1 and plasmid-borne Ldb16-FLAG with a photoreactive Bpa at the indicated positions were subjected to UV irradiation. Non-irradiated cells were used as controls. Solubilized membranes were subjected to immunoprecipitation with anti-FLAG antibodies, and bound proteins were analyzed by immunoblotting with FLAG, V5 and HA antibodies.
C) Lysates form Figure 1F that were not subjected to UV radiation. Sei1Δ cells expressing endogenously HA-tagged Ldo, and plasmid-borne Sei1-FLAG with a photoreactive Bpa at the indicated positions were not subjected to UV irradiation. Solubilized membranes were subjected to immunoprecipitation with anti-FLAG antibodies, and bound proteins were analyzed by immunoblotting with FLAG and HA antibodies.
D) Sei1Δ cells expressing endogenously HA-tagged Ldo and plasmid-borne Sei1-FLAG with a photoreactive Bpa at the indicated positions were subjected to UV irradiation. Non-irradiated cells were used as controls. Solubilized membranes were subjected to immunoprecipitation with anti-HA antibodies, and bound proteins were analyzed by immunoblotting with an HA antibody.

**Table S1 - List of strains used in this study**

| Strain | Identifier | Genotype | Origin |
|---|---|---|---|
| **BY4741** | yPC1505 | MATa *his3Δ1 leu2Δ0 met15Δ0 ura3Δ0* | (Brachmann et al. 1998)) |
| **FY251** | yPC1507 | MATa *his3Δ1 leu2Δ0 trp1Δ63 ura3-52* | Fred Winston |
| **BY4741** | yPC4288 | MATa *his3Δ1 leu2Δ0 met15Δ0 ura3Δ0, OSW5-3HA-HIS* | (Teixeira et al. 2018) |
| **BY4741** | yPC7472 | MATa *his3D1, leu2D0, met15D0, ura3D0m, Ldb16 1-133FLAG-HYGR OSW5-3HA-HIS* | This study |
| **BY4741** | yPC7474 | MATa *his3D1, leu2D0, met15D0, ura3D0m, Ldb16-FLAG-HYGR, OSW5-3HA-HIS* | This study |
| **FY251** | yPC12459 | MATa his3D200, *leu2D1, trp1D63, ura3-52, Ldb16::NAT,OSW5-3HA-KANR, fld1-V5-HYGB* | This study |
| **FY251** | yPC12630 | MATa *his3D200, leu2D1, trp1D63, ura3-52, Ldb16::NAT,OSW5-HA-KANR, fld1-V5-HYGB, Lro1::CRISPR, Are1::CRISPR, Are2::CRISPR, Dga1::CRISPR* | This study |
| **BY4741** | yPC12817 | MATa *his3D1, leu2D0, met15D0, ura3D0m, Ldb16 Δ44-59-CRISPR - FLAG-HYGR, OSW5-3HA-HIS* | This study |
| **BY4741** | yPC12818 | MATa *his3D1, leu2D0, met15D0, ura3D0m, Ldb16 S53,55,62-Ala T52,61,63 - Ala-CRISPR - FLAG-HYGR, OSW5-3HA-HIS* | This study |
| **BY4741** | yPC12853 | MATa *his3D1, leu2D0, met15D0, ura3D0m, Ldb16 1-133 Δ44-59-CRISPR - FLAG-HYGR, OSW5-3HA-HI* | This study |

| | | | |
|---|---|---|---|
| **BY4741** | yPC12854 | MATa *his3D1, leu2D0, met15D0, ura3D0m, Ldb16 1-133 S53,55,62-Ala T52,61,63 -Ala-CRISPR - FLAG-HYGR, OSW5-3HA-HIS* | This study |

**Table S2 - List of plasmids used in this study**

| Identifier | Plasmid | Origin |
|---|---|---|
| **bPC220** | pFA6a-3HA-kanMX6 | (Longtine et al. 1998) |
| **bPC557** | pESC-TRP *TyrRS-tRNA CUA* for pBpa incorporation | (Chin et al. 2003)) |
| **bPC720** | pFA6a-6xGLY-V5-hphMX4 | (Funakoshi and Hochstrasser 2009) |
| **bPC1125** | pRS415 $P_{gal}$*Dga1* | (Renne et al. 2022) |
| **bPC1126** | pRS415 $P_{gal}$*Are1* | (Renne et al. 2022) |
| **bPC1127** | pRS415 $P_{gal}$Lro1 | (Renne et al. 2022) |
| **bPC1940** | pRS416 $P_{ADH1}$ *Sei1-3xFLAG* $T_{ADH1}$ | (Klug et al. 2021) |
| **bPC2088** | pRS416 $P_{ADH1}$ *Sei1-L162 amber (TAG)-3xFLAG* $T_{ADH1}$ | (Klug et al. 2021) |
| **bPC2089** | pRS416 $P_{ADH1}$ *Sei1-T163 amber (TAG)-3xFLAG* $T_{ADH1}$ | (Klug et al. 2021) |
| **bPC2090** | pRS416 $P_{ADH1}$ *Sei1-S165 amber (TAG)-3xFLAG* $T_{ADH1}$ | (Klug et al. 2021) |
| **bPC2091** | pRS416 $P_{ADH1}$ *Sei1-P168 amber (TAG)-3xFLAG* $T_{ADH1}$ | (Klug et al. 2021) |
| **bPC2092** | pRS416 $P_{ADH1}$ *Sei1-Q169 amber (TAG)-3xFLAG* $T_{ADH1}$ | (Klug et al. 2021) |
| **bPC2093** | pRS416 $P_{ADH1}$ *Sei1-E170 amber (TAG)-3xFLAG* $T_{ADH1}$ | (Klug et al. 2021) |
| **bPC2094** | pRS416 $P_{ADH1}$ *Sei1-E172 amber (TAG)-3xFLAG* $T_{ADH1}$ | (Klug et al. 2021) |
| **bPC2095** | pRS416 $P_{ADH1}$ *Sei1-P176 amber (TAG)-3xFLAG* $T_{ADH1}$ | (Klug et al. 2021) |
| **bPC2096** | pRS416 $P_{ADH1}$ *Sei1-L179 amber (TAG)-3xFLAG* $T_{ADH1}$ | (Klug et al. 2021) |
| **bPC2097** | pRS416 $P_{ADH1}$ *Sei1-W186 amber (TAG)-3xFLAG* $T_{ADH1}$ | (Klug et al. 2021) |
| **bPC2145** | pRS416 $P_{ADH1}$ *Ldb16-3xFLAG* $T_{ADH1}$ | This study |

| | | |
|---|---|---|
| **bPC2146** | pRS416 $P_{ADH1}$*Ldb16 F30 amber (TAG)-3xFLAG* $T_{ADH1}$ | This study |
| **bPC2147** | pRS416 $P_{ADH1}$*Ldb16 F35 amber (TAG)-3xFLAG* $T_{ADH1}$ | This study |
| **bPC2148** | pRS416 $P_{ADH1}$*Ldb16 F49 amber (TAG)-3xFLAG* $T_{ADH1}$ | This study |
| **bPC2149** | pRS416 $P_{ADH1}$*Ldb16 S55 amber (TAG)-3xFLAG* $T_{ADH1}$ | This study |
| **bPC2150** | pRS416 $P_{ADH1}$*Ldb16 R56 amber (TAG)-3xFLAG* $T_{ADH1}$ | This study |
| **bPC2151** | pRS416 $P_{ADH1}$*Ldb16 V58 amber (TAG)-3xFLAG* $T_{ADH1}$ | This study |
| **bPC2152** | pRS416 $P_{ADH1}$*Ldb16 Q60 amber (TAG)-3xFLAG* $T_{ADH1}$ | This study |
| **bPC2153** | pRS416 $P_{ADH1}$*Ldb16 L64 amber (TAG)-3xFLAG* $T_{ADH1}$ | This study |
| **bPC2154** | pRS416 $P_{ADH1}$*Ldb16 F66 amber (TAG)-3xFLAG* $T_{ADH1}$ | This study |
| **bPC2155** | pRS416 $P_{ADH1}$*Ldb16 F76 amber (TAG)-3xFLAG* $T_{ADH1}$ | This study |
| **bPC2156** | pRS416 $P_{ADH1}$*Ldb16 F90 amber (TAG)-3xFLAG* $T_{ADH1}$ | This study |
| **bPC2161** | pRS416 $P_{ADH1}$*Ldb16 N33 amber (TAG)-3xFLAG* $T_{ADH1}$ | This study |
| **bPC2162** | pRS416 $P_{ADH1}$*Ldb16 L40 amber (TAG)-3xFLAG* $T_{ADH1}$ | This study |
| **bPC2163** | pRS416 $P_{ADH1}$*Ldb16 I42 amber (TAG)-3xFLAG* $T_{ADH1}$ | This study |
| **bPC2164** | pRS416 $P_{ADH1}$*Ldb16 I44 amber (TAG)-3xFLAG* $T_{ADH1}$ | This study |
| **bPC2165** | pRS416 $P_{ADH1}$*Ldb16 L54 amber(TAG)-3xFLAG* $T_{ADH1}$ | This study |
| **bPC2166** | pRS416 $P_{ADH1}$*Ldb16 L72 amber(TAG)-3xFLAG* $T_{ADH1}$ | This study |
| **bPC2167** | pRS416 $P_{ADH1}$*Ldb16 F73 amber(TAG)-3xFLAG* $T_{ADH1}$ | This study |
| **bPC2168** | pRS416 $P_{ADH1}$*Ldb16 V78 amber(TAG)-3xFLAG* $T_{ADH1}$ | This study |
| **bPC2169** | pRS416 $P_{ADH1}$*Ldb16 L79 amber(TAG)-3xFLAG* $T_{ADH1}$ | This study |

| | | |
|---|---|---|
| **bPC2170** | pRS416 *$P_{ADH1}$Ldb16 C80 amber(TAG)-3xFLAG $T_{ADH1}$* | This study |
| **bPC2571** | pRS416 *$P_{LDB16}$ Ldb16-3Xflag $T_{ADH1}$* | This study |

Table S3. Oligonucleotides used in this study.

| Primer | Nucleotide sequence (5′–3′) | Purpose |
|---|---|---|
| **5** | GCACGTCAAGACTGTCAAGG | For verification of strains tagged or deleted with pringle plasmid-based cassettes |
| **185** | CTATTGTACTCGAGCGAGGCAAGCTAAACAGATC | Reverse Primer 50 bp downstream ADH1 terminator |
| **488** | CATTTTTGTTAGAAAGGGTCAGGAAAAATCCAAGAAACATAGCggtcgacggatccccgggtt | For C-terminal tagging of Fld1 using Pringle-derived plasmids |
| **492** | TCTATCATTCACTTGTTAGTGCATGAGAAGAAGTAATTGCTCGATGAATTCGAGCTCGTT | Reverse primer for tagging/deletion of LDB16 ORF |
| **514** | GAGACTACTGCTAATAAAGCGGGTAATAAGTTCCAGCTCTCTggtcgacggatccccgggtt | For C-terminal tagging of Ldo16/45 using Pringle-derived plasmids |
| **515** | GACCTGCTAAACTTGCGAAAAATGTTTTTTTATTGCCGAGGtcgatgaattcgagctcgtt | Reverse primer for tagging/deletion of Ldo16/45 |
| **516** | CAACTACAGGAACCACAGGAG | F primer approx 200 5' of stop to verify C-terminal tags of Ldo16/45 |
| **570** | GCAACTGTAGGAGGAGAAAGCAGGTATATAACTAGCCGCAATCGGATCCCCGGGTTAATTAA | Forward primer for deletion of LDB16 ORF |
| **722** | TGCAGATTCAAAAGCTCCAA | F primer to verify deletion; ~500bp 5' ATG of Are1 |

| | | |
|---|---|---|
| **723** | GTCACAACACCTATAATCAT | R primer to verify deletion; ~500bp 3' stop of Are1 |
| **725** | TGCAGATTCAAAAGCTCCAA | F primer to verify deletion; ~500bp 5' ATG of Are2 |
| **726** | GTCACAACACCTATAATCAT | R primer to verify deletion; ~500bp 3' stop of Are2 |
| **727** | TGCAGATTCAAAAGCTCCAA | F primer to verify deletion; ~500bp 5' ATG of Lro1 |
| **729** | TGCAGATTCAAAAGCTCCAA | R primer to verify deletion; ~500bp 3' stop of Lro1 |
| **731** | TATAACAAGACGGAAAGATTGA | F primer to verify deletion; ~500bp 5' ATG of Dga1 |
| **732** | AAACCACCGGTATTTACTGC | R primer to verify deletion; ~500bp 3' stop of Dga1 |
| **763** | CTTAATGATTGCGGCCGCTGATAGACAACCACACGGTC | Forward Primer 450 bp upstream of LDB16 ORF |
| **3720** | AGGGGACGAAAATTAGCCGCTATTAATTCT | Fw primer 100 bp upstream of Are2 |
| **3721** | AGACGCTTGTAAATAATAAAAAAAA | Rv primer 100 bp downstream the stop codon of Are2 |
| **3722** | GAAGGACGAGCGTGGTGGAGGAAAGGGGCG | Fw primer 100 bp upstream of Are1 |
| **3723** | AGCTCTTTGCCCCTGTCTTTGACCACGGTG | Rv primer 100 bp downstream the stop codon of Are1 |
| **3724** | GAAAAATAAGAAATCTACTAAATACCGATA | Fw primer 100 bp upstream of Lro1 |
| **3725** | TTTTGTTTTTGTTTTCTGTTTATTTTCTAG | Rv primer 100 bp downstream the stop codon o Lro1 |
| **3726** | GTTCCATTAAGGAGGTTTACTATCATCTCA | Fw primer 100 bp upstream of Dga1 |

| | | |
|---|---|---|
| **3727** | TATTCCTGTAAGTTAATACTCTTACTTAAG | Rv primer 100 bp downstream the stop codon of Dga1 |
| **3843** | GTTGACAAAGACCAAGACCA | 100 nmol Duplex with gRNA sequence GTTGACAAAGACCAAGACCA with restriction sites for ligation in pML107. For deletion of Are1 |
| **3844** | cttcaccgtagtcttcaccg | 100 nmol Duplex with gRNA sequence cttcaccgtagtcttcaccg with restriction sites for ligation in pML107. For deletion of Are1 |
| **3845** | GCAGGAGGAGGAGTACCCTG | 100 nmol Duplex with gRNA sequence GCAGGAGGAGGAGTACCCTG with restriction sites for ligation in pML107. For deletion of Are1 |
| **3846** | AATTACACAGAATCACAGCG | 100 nmol Duplex with gRNA sequence AATTACACAGAATCACAGCG with restriction sites for ligation in pML107. For deletion of Are2 |
| **3847** | ACAATACTTATGCAAGTGGG | 100 nmol Duplex with gRNA sequence ACAATACTTATGCAAGTGGG with restriction sites for ligation in pML107. For deletion of Are2 |
| **3848** | CAAAAGACAATCTATAACAG | 100 nmol Duplex with gRNA sequence CAAAAGACAATCTATAACAG with restriction sites for ligation in |

| | | |
|---|---|---|
| | | pML107. For deletion of Are2 |
| **3849** | AGTGCAAAAAGAAATGAGCG | 100 nmol Duplex with gRNA sequence AGTGCAAAAAGAAATGAGCG with restriction sites for ligation in pML107. For deletion of Lro1 |
| **3850** | GACAGGAAAAGAGACGGGAA | 100 nmol Duplex with gRNA sequence GACAGGAAAAGAGACGGGAA with restriction sites for ligation in pML107. For deletion of Lro1 |
| **3851** | ATGTGTCACAAATGGGCCCA | 100 nmol Duplex with gRNA sequence ATGTGTCACAAATGGGCCCA with restriction sites for ligation in pML107. For deletion of Lro1 |
| **3852** | gaagaaggaagCCCTACAGC | 100 nmol Duplex with gRNA sequence gaagaaggaagCCCTACAGC with restriction sites for ligation in pML107. For deletion of Dga1 |
| **3853** | CATTCAATGATATaagaaga | 100 nmol Duplex with gRNA sequence CATTCAATGATATaagaaga with restriction sites for ligation in pML107. For deletion of Dga1 |
| **3854** | TTTCCATGATTTGTATATTG | 100 nmol Duplex with gRNA sequence TTTCCATGATTTGTATATTG with restriction sites for ligation in |

| | | |
|---|---|---|
| | | pML107. For deletion of Dga1 |
| **3910** | GAAGGACGAGCGTGGTGGAGGAAAGGG GCGCCATTGGCACACTCACGCAGGTGGT TGTTCAGCACGGCTTGCAGCAAGAGCGC CAAAACAGATTGCAAGAACTGCCACCAT ACCACGTGTGTCCCTCGCAAGCCCTTGA TAGATATACAATAGGGAATGGGCGTCC GCTCCACCGTGGTCAAAGACAGGGGCAA AGAGCT | 4 nmole single strand oligo of 100 bp upstream the Are1 ORF plus 100 bp downstream the stop codon. |
| **3911** | AGGGGACGAAAATTAGCCGCTATTAAT TCTGGTATTGCCACCTAGACAAGAAGTA AACAGACACATTACGTTAGCAAAAGCA ACAATAACAAACACAACCGGCATCCTGC AACTGTTCTGTGGAGCTATTAAATCTTT ATAGTAAATTTTTTTTTACTTTTTTTTT TTTTT TTTTTTTTTTTTATTATTTACAAGCGTC T | 4 nmole single strand oligo of 100 bp upstream the Are2 ORF plus 100 bp downstream the stop codon. Used for deletion of the ORF by CRISPR. |
| **3912** | GAAAAATAAGAAATCTACTAAATACCG ATACGAAGAAGCGTATAGTAACAGCCA TTACAAAAGGTTCTCTACCAACGAATTC GGCGACAATCGAGTAAAACTCACTATCC ATCCGTGTATTATTTCAAAGAGCGAAA AGAAGGCGCGTCGCGTCGACGCGCCTTT TTAGGCTAGAAAATAAACAGAAAACAA AAACAAAA | 4 nmole single strand oligo of 100 bp upstream the Lro1 ORF plus 100 bp downstream the stop codon. Used for deletion of the ORF by CRISPR. |
| **3914** | GTTCCATTAAGGAGGTTTACTATCATCT CATTTCCATTTACACATACACTTACATA TACATAAGGAAACGCAGAGGCATACAG TTTGAACAGTCACATAATAATGAATTC ATTGGAAAACACAAAATATGTTAGAAT AAATAAGGATTTTTTAGTGTTTGGGCT GTATATCTTAAGTAAGAGTATTAACTT ACAGGAATA | gBlock of 100 bp upstream the Dga1 ORF plus 100 bp downstream the stop codon. Used for deletion of the ORF by CRISPR. |
| **4195** | CAATTTTCCATTGCATAATATGTGTACT TTAGTTTATCACAGGTTGCACTGCATTC | Generating amber codon site in Sei1 F263 Forward |
| **4196** | GAATGCAGTGCAACCTGTGATAAACTA AAGTACACATATTATGCAATGGAAAAT TG | Generating amber codon site in Sei1 F263 Reverse |

| | | |
|---|---|---|
| **4197** | ACGGATTCCATGTCGCCTCAGTAGATCG AACAACTAGGCCCATCACGTCTAG | Generating amber codon site in Sei1 E170 Forward |
| **4198** | TGATGGGCCTAGTTGTTCGATCTACTGA GGCGACATGGAATCCGTCAGTGCG | Generating amber codon site in Sei1 E170 Reverse |
| **4199** | GATTCCATGTCGCCTCAGGAGATCTAGC AACTAGGCCCATCACGTCTAGAC | Generating amber codon site in Sei1 E172 Forward |
| **4200** | GTCTAGACGTGATGGGCCTAGTTGCTAG ATCTCCTGAGGCGACATGGAATC | Generating amber codon site in Sei1 E172 Reverse |
| **4219** | CTATTGTCTGCCTCGCATAGACGGATTC CATGTCGCCTC | Generating amber codon site in Sei1 L162 Forward |
| **4220** | GACATGGAATCCGTCTATGCGAGGCAGA CAATAGGTCTAG | Generating amber codon site in Sei1 L162 Reverse |
| **4221** | GATCGAACAACTAGGCCCATCACGTTAG GACGTTTACGATGAAGAATGGC | Generating amber codon site in Sei1 L179 Forward |
| **4222** | GCCATTCTTCATCGTAAACGTCCTAACG TGATGGGCCTAGTTGTTCGATC | Generating amber codon site in Sei1 L179 Reverse |
| **4227** | CTCGCACTGACGGATTCCATGTCGTAGC AGGAGATCGAACAACTAGGCCCATCAC | Generating amber codon site in Sei1 P168 Forward |
| **4228** | GTGATGGGCCTAGTTGTTCGATCTCCTG CTACGACATGGAATCCGTCAGTGCGAG | Generating amber codon site in Sei1 P168 Reverse |
| **4229** | CAGGAGATCGAACAACTAGGCTAGTCAC GTCTAGACGTTTACGATG | Generating amber codon site in Sei1 P176 Forward |
| **4230** | CATCGTAAACGTCTAGACGTGACTAGCC TAGTTGTTCGATCTCCTG | Generating amber codon site in Sei1 P176 Reverse |
| **4231** | ACTGACGGATTCCATGTCGCCTTAGGAG ATCGAACAACTAGGCCCATCACGTC | Generating amber codon site in Sei1 Q169 Forward |

| | | |
|---|---|---|
| **4232** | GACGTGATGGGCCTAGTTGTTCGATCTC CTAAGGCGACATGGAATCCGTCA | Generating amber codon site in Sei1 Q169 Reverse |
| **4233** | ACTGACGGATTAGATGTCGCCTCAGGAG ATCGAAC | Generating amber codon site in Sei1 S165 Forward |
| **4234** | CTCCTGAGGCGACATCTAATCCGTCAGT GCGAGGCAGAC | Generating amber codon site in Sei1 S165 Reverse |
| **4235** | GTCTGCCTCGCACTGTAGGATTCCATGT CGCCTCAGGAG | Generating amber codon site in Sei1 T163 Forward |
| **4236** | AGGCGACATGGAATCCTACAGTGCGAG GCAGACAATAGGTCTAG | Generating amber codon site in Sei1 T163 Reverse |
| **4237** | CTAGACGTTTACGATGAAGAATAGCTA AATACAATAAGAATAGAGG | Generating amber codon site in Sei1 W186 Forward |
| **4238** | CCTCTATTCTTATTGTATTTAGCTATTC TTCATCGTAAACGTCTAGACG | Generating amber codon site in Sei1 W186 Reverse |
| **4239** | CTTATTTAGAAGTGGCGCGCCTCACTTG TCATCGTCATCCTTGTAG | Generating Sei1 tagging with 3xFLAG in pRS416 |
| **5289** | CAGCTAGGTTTTAAAATTATATAGCGA GAAGTACAATTtcgatgaattcgagctcgtt | For C-terminal tagging of Fld1 with either FLAG or V5 from bPC 713 and 720 respectively. |
| **5290** | GAAGTGGCGCGCCtcACGTAGAATCGAG ACCGAGGAG | For swapping the stop codon on the V5 TAG |
| **5291** | TATCACGAGGCCCTTTCGTCTC | For verification of V5 tagged strains |
| **5292** | GCACATTAGACTagTATGTGGTTTTGAC GTTGTTTCAATATTTTG | For introducing an amber stop codon at F66 into an Ldb16 containing plasmid |
| **5293** | TCAAAACCACATActAGTCTAATGTGCT TGTTTGTGCAACTAAAC | For introducing an amber stop codon at F66 into an Ldb16 containing plasmid |

| | | |
|---|---|---|
| **5294** | GTTTTGACGTTGTTTCAATATTagGCTGTGCTTTGTGCTTTTGGC | For introducing an amber stop codon at F76 into an Ldb16 containing plasmid |
| **5295** | GCACAGCctAATATTGAAACAACGTCAAAACCACATAGAAGTC | For introducing an amber stop codon at F76 into an Ldb16 containing plasmid |
| **5296** | CAATCAACTTCCGGAGTGTAAAAACTGATTTTCAATGTTTGTGGTGGATTGGAGC | For amplification of Ldb16 from pringle plasmids |
| **5297** | TTTAGTTGCAtAgACAAGCACATTAGACTTCTATGTGG | For introducing an amber stop codon at Q60 into an Ldb16 containing plasmid |
| **5298** | ATGTGCTTGTcTaTGCAACTAAACGGCTCAGCGACG | For introducing an amber stop codon at Q60 into an Ldb16 containing plasmid |
| **5299** | TAATGTGCTTGTTTGTGCAACTAActaGCTCAGCGACGTCCCCAGGAACAAC | For introducing an amber stop codon at R56 into an Ldb16 containing plasmid |
| **5300** | CGTCGCTGAGCtagTTAGTTGCACAAACAAGCACATTAGAC | For introducing an amber stop codon at R56 into an Ldb16 containing plasmid |
| **5301** | CGTCGCTGtagCGTTTAGTTGCACAAACAAGCACATTAGAC | For introducing an amber stop codon at S55 into an Ldb16 containing plasmid |
| **5302** | CAACTAAACGctaCAGCGACGTCCCCAGGAACAAC | For introducing an amber stop codon at S55 into an Ldb16 containing plasmid |
| **5313** | TGCCTTTATCAATTTTCGTATGGTAGGGGTTTCAGCTTGTGGCATTG | For introducing an amber stop codon at N33 into an Ldb16 containing plasmid |
| **5314** | GAAACCCCTACCATACGAAAATTGATAAAGGCAGTTGC | For introducing an amber stop codon at |

| | | |
|---|---|---|
| | | N33 into an Ldb16 containing plasmid |
| **5315** | TTTCAGCTTGTGGCATaGCCTATCAACATCCCATTGAG | For introducing an amber stop codon at L40 into an Ldb16 containing plasmid |
| **5316** | ATGTTGATAGGCtATGCCACAAGCTGAAACCCATTC | For introducing an amber stop codon at L40 into an Ldb16 containing plasmid |
| **5317** | TTGTGGCATTGCCTtagAACATCCCATTGAGGTTGTTCCTGGG | For introducing an amber stop codon at I42 into an Ldb16 containing plasmid |
| **5318** | CTCAATGGGATGTTctaAGGCAATGCCACAAGCTGAAACC | For introducing an amber stop codon at I42 into an Ldb16 containing plasmid |
| **5319** | TTGTGGCATTGCCTATCAACtagCCATTGAGGTTGTTCCTGGG | For introducing an amber stop codon at I44 into an Ldb16 containing plasmid |
| **5320** | CTCAATGGctaGTTGATAGGCAATGCCACAAGCTGAAACC | For introducing an amber stop codon at I44 into an Ldb16 containing plasmid |
| **5321** | TGGGGACGTCGtaGAGCCGTTTAGTTGCACAAACAAG | For introducing an amber stop codon at L54 into an Ldb16 containing plasmid |
| **5322** | AACTAAACGGCTCtaCGACGTCCCCAGGAACAACCTC | For introducing an amber stop codon at L54 into an Ldb16 containing plasmid |
| **5323** | TTCTATGTGGTTTTGACGTaGTTTCAATATTTTGCTGTGCTTTG | For introducing an amber stop codon at L72 into an Ldb16 containing plasmid |
| **5324** | AGCAAAATATTGAAACtACGTCAAAACCACATAGAAGTCTAATG | For introducing an amber stop codon at L72 into an Ldb16 containing plasmid |

| | | |
|---|---|---|
| **5325** | GTGGTTTTGACGTTGTagCAATATTTTGCTGTGCTTTGTGCTTTT | For introducing an amber stop codon at F73 into an Ldb16 containing plasmid |
| **5326** | AGCAAAATATTGctACAACGTCAAAACCACATAGAAGTCTAATGTG | For introducing an amber stop codon at F73 into an Ldb16 containing plasmid |
| **5327** | TCAATATTTTGCTtaGCTTTGTGCTTTTGGCAGCATCATAGGAC | For introducing an amber stop codon at V78 into an Ldb16 containing plasmid |
| **5328** | AAAAGCACAAAGCtaAGCAAAATATTGAAACAACGTCAAAACCAC | For introducing an amber stop codon at V78 into an Ldb16 containing plasmid |
| **5329** | TCAATATTTTGCTGTGtagTGTGCTTTTGGCAGCATCATAGGAC | For introducing an amber stop codon at L79 into an Ldb16 containing plasmid |
| **5330** | AAAAGCACActaCACAGCAAAATATTGAAACAACGTCAAAACCAC | For introducing an amber stop codon at L79 into an Ldb16 containing plasmid |
| **5331** | TCAATATTTTGCTGTGCTTTagGCTTTTGGCAGCATCATAGGACTCATC | For introducing an amber stop codon at C80 into an Ldb16 containing plasmid |
| **5332** | AAAAGCctAAAGCACAGCAAAATATTGAAACAACGTCAAAACCAC | For introducing an amber stop codon at S55 into an Ldb16 containing plasmid |
| **5349** | GTTAATTAACGGTTGAGGCGCGCCACTTCTAAATAAGC | To amplify the ADH terminator with a SgsI site |
| **5371** | taacccggggatccgtcgaccAATGTTGACAATACTAGTGGAAG | To truncate Ldb16 on pringle plasmids and add a SalI site at the 3' end |
| **5392** | GATCTCCCATTGAGGTTGTTCCTGGTTTTAGAGCTAG | 100 nmol Duplex with gRNA sequence with restriction sites for |

| | | |
|---|---|---|
| | | ligation in pML107. For mutating Ldb16. Guide 1 forward |
| **5393** | CTAGCTCTAAAACCAGGAACAACCTCAATGGGA | 100 nmol Duplex with gRNA sequence with restriction sites for ligation in pML107. For mutating Ldb16. Guide 1 reverse |
| **5394** | GATCAACGGCTCAGCGACGTCCCCGTTTTAGAGCTAG | 100 nmol Duplex with gRNA sequence with restriction sites for ligation in pML107. For mutating Ldb16. Guide 2 forward |
| **5395** | CTAGCTCTAAAACGGGGACGTCGCTGAGCCGTT | 100 nmol Duplex with gRNA sequence with restriction sites for ligation in pML107. For mutating Ldb16. Guide 2 reverse |
| **5396** | GATCTGCTTGTTTGTGCAACTAAAGTTTTAGAGCTAG | 100 nmol Duplex with gRNA sequence with restriction sites for ligation in pML107. For mutating Ldb16. Guide 3 forward |
| **5397** | CTAGCTCTAAAACTTTAGTTGCACAAACAAGCA | 100 nmol Duplex with gRNA sequence with restriction sites for ligation in pML107. For mutating Ldb16. Guide 3 reverse |
| **5398** | CACCTCTCTTCCGTGCCCTCGTGCAACTGCCTTTATCAATTTTCGTATGGAATGGGTTTCAGCTTGTGGCATTGCCTATCAACGCTGCCAGTGCGGCTTCTGCGGCCTCACAAACAAGCACATTAGACTTCTATGTGGTTTTGACGTTGTTTCAATATTTTGCTGTGCTTTGTGCTTTTGGCAGCATCATAGGACTCATC | template for creating the Δhelix mutation in Ldb16 and Ldb16 1-133 by CRISPR |

| | | |
|---|---|---|
| **5399** | TGCAACTGCCTTTATCAATTTTCGTATGGAATGGGTTTCAGCTTGTGGCATTGCCTATCAACATCCCATTGAGGTTGTTtCTGGGaGCTGCACTGGCACGTTTAGTTGCACAAGCTGCAGCGTTAGACTTCTATGTGGTTTTGACGTTGTTTCAATATTTTGCTGTGCTTTGTGCTTTTGGCAGCATCATAGGACTCATC | template for creating the 6A mutation in Ldb16 and Ldb16 1-133 by CRISPR |
| **5400** | CACCTCTCTTCCGTGCCCTCG | primer to amplify the CRISPR Δhelix template |
| **5401** | GATGAGTCCTATGATGCTGCCAAAAG | primer to amplify both CRISPR templates |
| **5402** | TGCAACTGCCTTTATCAATTTTCG | primer to amplify the CRISPR 6A template |
